# MeCP2 NID interaction with RNA: Implications for Rett Syndrome-Relevant Protein Regulation

**DOI:** 10.1101/2025.11.19.689340

**Authors:** Katrina V. Good, Hilmar Strickfaden, Tahir Muhammad, John B. Vincent, Michael Hendzel, Christopher J. Nelson, Juan Ausió

**Affiliations:** Department of Biochemistry and Microbiology, University of Victoria, Victoria, BC V8W 3P6, Canada; Molecular Neuropsychiatry & Development (MiND) Lab, Campbell Family Mental Health Research Institute, Cantre for Addiction and Mental Health, Toronto, ON M5T 1A8, Canada; Department of Oncology, Faculty of Medicine and Dentistry, University of Alberta, Edmonton, AB T6G 1Z2, Canada; Institute of Medical Science, University of Toronto, Toronto, ON M5S 1A8, Canada; Department of Psychiatry, University of Toronto, Toronto, ON M5T 1R8, Canada

**Keywords:** MeCP2, RNA Binding, Rett Syndrome, NCoR, NEAT1

## Abstract

Mutations in the *MECP2* gene cause the progressive neurodevelopmental disorder Rett syndrome. Pathogenic missense mutation hotspots exist in the protein’s Methyl DNA binding Domain (MBD), and the NCoR Interaction Domain (NID), indicating these regions as critical for MeCP2 function. The NID binds to a co-repressor complex allowing transcriptional repression at target genes. A putative RNA Binding Domain (RBD) was identified that overlaps with the NID, yet the role that RNA interaction plays in MeCP2 function remains underexplored. Using cell-based and *in vitro* molecular assays, we validated RNA interaction at the NID/RBD of MeCP2 both to a dsRNA probe *in vitro* and to the lncRNA *NEAT1_2* in cells. As expected, this region did not appear to affect MeCP2-chromatin interactions; however, we found that RNA-RBD interaction precludes MeCP2-NCoR binding in cells. Taken together, we find that RNA interaction at this non-canonical RNA binding domain regulates important MeCP2-protein interactions and therefore may be a key part of the pathophysiology of Rett syndrome.

## Introduction

Methyl CpG Binding Protein 2 (MeCP2) is a methyl CpG DNA binding protein with intrinsically disordered regions flanking its structured Methyl DNA Binding Domain (MBD), providing a flexible backbone to which over 40 known protein partners interact, eliciting function in processes like transcriptional regulation, miRNA biogenesis, chromatin remodeling, and mRNA splicing (1). Rett syndrome (RTT) (OMIM: 312750), caused by *de novo MECP2* mutations is associated not only with neurological dysfunction, but also respiratory, gastrointestinal, and inflammatory conditions due to effects downstream of altered brain function as well as direct changes to *MECP2* in those tissues, as it is expressed ubiquitously in other somatic cells (2, 3). The mechanism(s) responsible for regulating MeCP2’s disparate interactions, and therefore its multiplex functionality, remain unclear. Non-coding RNAs (ncRNAs) have quickly become established as important regulatory molecules, especially in nuclear processes (4).

Past and recent evidence exists for MeCP2-RNA binding both *in vitro* and *in vivo*. The *in vitro* data confirm direct dsRNA and not ssRNA binding, the more recent of which suggests preference for bulges and extended stem-loops (5, 6); however, the physiological relevance of the RNAs identified were not explored. MeCP2-RNA binding *in vivo* has been demonstrated to a variety of RNA species. In mouse primary cortical neurons, MeCP2 immunoprecipitates microRNAs (miRNAs) whose targets are dysregulated in *Mecp2*-null brain (7); perhaps not unrelated is MeCP2’s role in supressing miRNA processing through interaction with DiGeorge syndrome critical region 8 (DGCR8) (8). MeCP2 binding to circular RNA (circRNA) was shown to regulate sirtuin 2 (SIRT2) levels in human neural progenitor cells (9). Some reports suggest RNA-dependent interactions of MeCP2 with mRNA splicing factors (10, 11), although mRNA enrichment over the transcriptome was not seen in RNA immunoprecipitation followed by sequencing (RIP-seq) results in male mouse cerebellum (12). Instead, several long non-coding RNAs (lncRNAs) were enriched by MeCP2 RIP-seq, including Retinal non-coding RNA 3 (*Rncr3*), with enrichment in exons 2-3 and disrupted splicing in *Mecp2*-null mouse cerebellum, indicating a role in lncRNA splicing. Specific regulation by lncRNA-MeCP2 binding has also been shown wherein *NEAT1_2* transcript and MeCP2 protein levels are reciprocally regulated by MeCP2-RNA binding in Huntington disease patient-derived cells and *Evf2* (*DLX6_AS* in human) recruits MeCP2 to regulate its gene target in developing mouse brain (13, 14). MeCP2 is also regulated by interaction with ncRNAs outside of a neuronal context, such as in mouse heart tissue, where the lncRNA *Mhrt* recruits MeCP2 to the *Pri-miR-208b* promoter in a sex-dependent manner (15) and in human cancer cell lines in which aberrantly expressed high-copy satellite II (*HSATII*) RNA recruits MeCP2 into cancer-specific puncta which is associated with genomic instability and cell division defects (16, 17).

An RNA Binding Domain (RBD) was identified through RNA pull down and mass spectrometry which is described as non-canonical (18), meaning it lacks homology to known RBDs. Rather than canonical RBDs acting upon specific RNA molecules, it is suggested that RNA acts upon non-canonical RBDs to regulate protein function (19). Intriguingly, this RBD almost perfectly overlaps the NCoR Interaction Domain (NID) of MeCP2, which is responsible for recruiting the Nuclear receptor co-repressor (NCoR) complex – a critical facet of MeCP2 function – and is a hotspot for Rett syndrome-causing missense mutations, which are cumulatively responsible for approximately 6% of RTT cases (20).

Here we show that the RBD/NID contributes to MeCP2-RNA binding *in vitro* and *in vivo* and that this interaction regulates MeCP2 binding to co-repressor protein partners, adding to the repertoire of known mechanisms guiding MeCP2 protein regulation.

Through its many molecular interactions, MeCP2 serves as an important metabolic node wherein various inputs modulate protein behaviour and downstream effects. The findings herein are crucial to appreciate the full complexity of Rett syndrome progression.

## Results

### Preamble: A note on MeCP2 isoforms and nomenclature

Two MeCP2 protein isoforms exist which differ only in their N-terminal ends (**Fig. 1A**). MeCP2 E1 is expressed and binds to DNA more dynamically than the E2 isoform and is the primary isoform present in the brain (21, 22) whereas MeCP2 E2 is prominent in other cell types, suggesting differences that are critical for understanding RTT pathology. However, MeCP2 functions at the region of interest here are likely not affected by these N-terminal differences, as evidenced by binding shown to the NCoR co-repressor complex when these N-terminal regions are removed (23), as well as mutations in the NID having little to no effect on the chromatin localization of MeCP2, as discussed below. As such, it is worth noting that, in the present study, the exogenously expressed MeCP2 in cellular assays herein was the E1 type whereas the *in vitro* binding assays were performed with recombinantly expressed MeCP2 E2, because the E1 sequence was providing relatively low yields in our hands. Therefore, in the case of the lysine (K) to glutamic acid (E) missense mutant used in this work, the numbering of K304E and K316E refer to the same mutation in isoforms E2 and E1, respectively.

**Figure 1.**
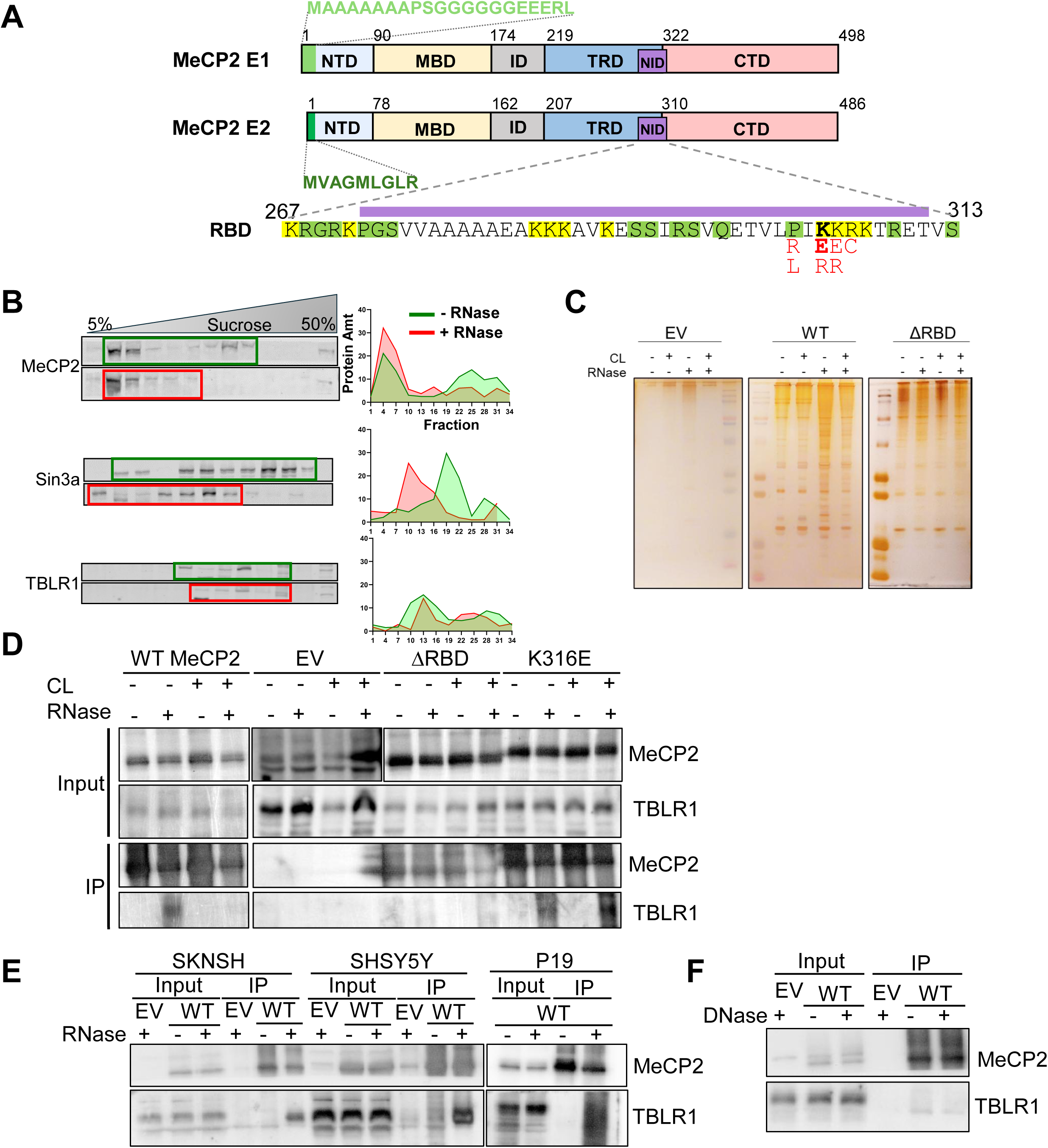
Important MeCP2-Protein Interactions are affected by MeCP2 RBD-RNA Binding. **A)** Schematic of MeCP2 protein domains, with differences between the two isoforms as well as the RNA Binding Domain and overlapping NCoR Interaction Domain shown, with K304 in bold and select RTT-causing missense mutations in red. In the RBD sequence, yellow and green highlights indicate K-rich and disorder-promoting regions, respectively. NTD: N-Terminal Domain, MBD: Methyl CpG DNA Binding Domain, ID: Intervening Domain, TRD: Transcriptional Repression Domain, NID: NCoR Interaction Domain, CTD: C-Terminal Domain, RBD: RNA Binding Domain. **B)** Western blots of proteins after sucrose gradient ultracentrifugation with and without RNase treatment of HeLa S3 cell nuclear lysates. Quantifications (right) are based on densitometric measurements of WB signals normalized to the total protein measured from all fractions and plotted using GraphPad Prism. Due to the complexity of the procedure and large sample volumes required, n=2 biological replicates with four technical replicate western blots were tested. **C)** Silver Stain of MeCP2-FLAG CoIP samples from HEK293 cells with and without UV-crosslinking and/or RNase treatment. Image represents 3 separate experiments. **D-F)** Co-IP and western blot of indicated proteins by transiently transfected Flag-tagged MeCP2 with indicated conditions. Images represent at least three separate experiments.

### Important MeCP2-Protein Interactions are affected by MeCP2 RBD-RNA Binding

We first set out to identify whether RNA interaction at the NID/RBD could regulate MeCP2 activities, starting with MeCP2-protein interactions. Given the unknown specificity of MeCP2 for RNA, research describing RNA-dependent protein complex formation using sucrose gradient ultracentrifugation prompted the first step (24). According to this approach, protein complexes that require RNA for their formation will disassemble upon RNase A treatment, leading to a redistribution of proteins to lower density sucrose after centrifugation, which is then fractionated and probed for proteins of interest (**Fig. S1A, B**). While MeCP2 was not detected by mass spectrometry in HeLa cells in the original article, known MeCP2 protein partners, Sin3a and NCoR, displayed RNA dependent and independent fractionation, respectively. Additionally, the sucrose gradient migration of other known MeCP2 interacting proteins showed varying RNA-dependencies (**Fig. S1C**) (24). Given these observations, we tested for an RNA-dependent shift of MeCP2 in HeLa S3 cells, using TBLR1 of the NCoR complex and Sin3a as negative and positive controls for protein shift (**Fig. 1B**). Without RNase treatment, MeCP2 is detectable across most of the gradient, suggesting multiple MeCP2-protein complex species. After RNase A treatment, MeCP2 shifts away from higher density fractions, suggesting that some MeCP2-protein complex interactions depend on RNA. In support of this interpretation, silver stained SDS-PAGE gels of proteins co-immunoprecipitated (Co-IP) by MeCP2-FLAG transfected into HEK293 cells show an RNA-dependent change to the MeCP2-protein interactome as a function of the RBD (**Fig. 1C**). The abundance of immunoprecipitated protein partners of MeCP2^WT^ increases after RNase treatment, whereas that of the RBD deletion mutant (MeCP2^ΔRBD^) was relatively unchanged, implicating this region in both MeCP2-RNA and MeCP2-protein interactions. We also employed a UV-crosslinking step, which creates covalent crosslinks between RNA and protein that are at a zero distance (i.e. bound) at the time of crosslinking (25, 26). Crosslinking alone does not appear to impact bulk MeCP2^WT^ interactions, and RNase treatment with crosslinking only marginally affect the resulting interactome compared to RNase alone. MeCP2^ΔRBD^ appears to render RNase A and crosslinking ineffective indicating that the increased MeCP2^WT^ protein interactions upon RNase treatment occurs at the NID/RBD. Thus, MeCP2-RNA interaction at the NID/RBD prevents many MeCP2-protein interactions, while its involvement in large macromolecular complexes may depend on the presence of RNA.

Given the overlapping NID and RBD and the reported importance of NCoR complex binding to MeCP2 function and RTT pathology, we tested how this interaction is affected by RNA binding. It is well established that the direct NCoR complex binding partner to MeCP2, TBLR1, interacts at the NID (23, 27). That TBLR1 was only immunoprecipitated by MeCP2-FLAG in RNase treated HEK293 nuclei (**Fig. 1D**) indicates that TBLR1-MeCP2 interaction is negatively regulated by RNA at the NID/RBD. We hypothesized that because UV-crosslinking creates covalent RNA-protein links that are at a zero distance at the time of crosslinking, RNA-MeCP2 crosslinks, if binding occurs at the NID, would be inaccessible to RNase and obstruct MeCP2^WT^-TBLR1 binding at the NID even when RNase is applied to the sample. This was observed, supporting direct RNA binding at the NID region of MeCP2 *in vivo*.

Expectedly, RBD deletion precludes TBLR1 binding. These data are in line with our silver stain results displaying a lack of increased MeCP2^WT^ interactions with crosslinked RNase treated samples, and a similar lack of increase under any condition by MeCP2^ΔRBD^, indicating the NID/RBD facilitates both RNA and protein interactions.

An RTT-causing mutation was previously shown to negatively impact MeCP2-TBLR1 binding (27), MeCP2^K316E^, immunoprecipitates TBLR1 with RNase treatment, regardless of crosslinking. This may be because the mutation, while not substantial enough to completely abrogate TBLR1 binding, impacts MeCP2 crosslinking to RNA rendering it ineffective, allowing protein-protein interaction; or, perhaps TBLR1 binds despite crosslinking due to RNA interaction occurring primarily at other residues within the NID. We confirmed RNase-dependent TBLR1 immunoprecipitation in other cell lines representing mouse and human neuronal cells (**Fig. 1E**) given the reported RTT-relevance to MeCP2-TBLR1 interaction, as well as protein atlas data suggesting relatively low *TBL1XR1* mRNA levels in HEK293 cells (**Fig. S2A**) (28) (proteinatlas.org). DNase treatment did not allow MeCP2-TBLR1 Co-IP (**Fig. 1F**), confirming that this is an RNA-specific phenomenon, whereas past MeCP2-TBLR1 IP assays customarily included the DNA and RNA endonuclease Benzonaze prior to Co-IP (27, 29). MeCP2 binds to another HDAC-containing co-repressor complex, Sin3a, through residues 162-279 (174-191 in E1), upstream of TBLR1-interacting residues (30) . Wild type and mutant MeCP2 immunoprecipitated Sin3a under all conditions tested, but the IP signal was strongest in the RNase treatment and crosslinking condition (**Fig. S2B**), which we hypothesize may be due to loss of physical occlusion by either RNA or other protein complexes, like NCoR, modeled in **Figure 5**. Altogether, our data thus far suggest both direct and indirect regulation of MeCP2 RBD-protein interactions by RNA.

### MeCP2-chromatin distribution and affinity are unaffected by RBD mutations or RNase treatment

MeCP2-chromatin interaction is another important broad category of MeCP2 function that was important to evaluate in relation to RNA binding. Confocal microscopy imaging showed unchanged Pearson’s correlation coefficient (PCC) between MeCP2-GFP and DAPI-dense signal upon RBD mutation in C2C12 mouse myoblast cells (**Fig. 2A**), which were chosen based on the presence of AT-rich chromocenters in their nuclei to which MeCP2 is known to localize and for which DAPI has preference (31). Our data is corroborated by previous works showing mutations in this region of the protein having little to no effect on MeCP2-chromatin association (32, 33). MeCP2 chromatin distribution analysis via Micrococcal Nuclease (MNase) digestion is well established, wherein the majority of MeCP2 associates with the most MNase accessible fraction (S1) (34). While transient over-expression of MeCP2 obscures the expected MeCP2-chromatin distribution, MeCP2^ΔRBD^ chromatin distribution is unchanged compared to the wild type counterpart (**Fig. 2B**). Endogenous MeCP2-chromatin distribution is also similar between adult postnatal day 30 mouse whole brain and liver lysates (**Fig. 2C**). Liver tissue is known to be abundant in RNAses (35); thus, presumably any role played by RNA in global MeCP2-chromatin distribution would result in different chromatin distributions between the two tissues upon cell and nuclear lysis in the experimental procedure. Additionally, Salt Extraction of endogenous MeCP2 from chromatin is unchanged in HEK293 cells with or without RNase treatment (**Fig. 2D**), indicating that RNA interaction with MeCP2 does not change its affinity for chromatin in cells, whereas DNA methylation levels were previously shown to impact MeCP2 release from chromatin (34). NID/RBD mutations did affect MeCP2 kinetics in NIH/3T3 mouse fibroblast nuclei (which have AT-rich chromocenters), as measured through fluorescence recovery after photobleaching (FRAP) (**Fig. 2E, Fig. S3**). However, the difference is marginal compared to that seen with the MBD-disrupting T158M missense mutation. Altogether, these data rule out a significant role for MeCP2 RBD-RNA binding in global MeCP2-chromatin interaction in cells.

**Figure 2.**
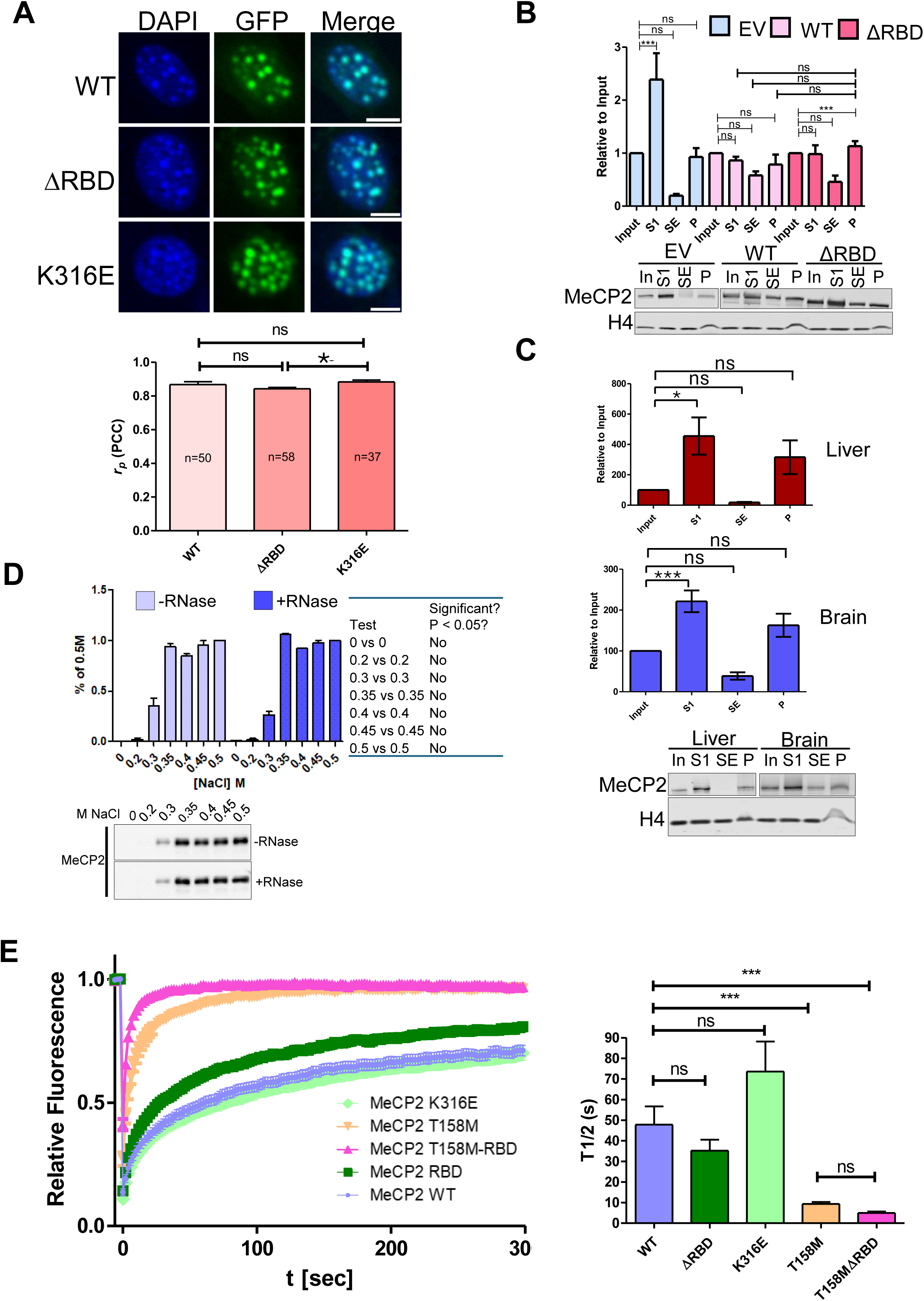
MeCP2-chromatin distribution and affinity are unaffected by RBD mutations or RNase treatment. **A)** (top) Representative confocal microscopy images of C2C12 cells transiently expressing MeCP2-GFP; scale bar = 10 μM. (bottom) Pearson’s Correlation Coefficient analysis between DAPI and GFP signal. Data represent number of nuclei indicated and three separate transfections. **B and C)** Micrococcal Nuclease digestion-based separation of chromatin into its most (S1) to least (P) nuclease accessible fractions and their associated components. N=3, 3 technical replicates each. **D)** Salt (NaCl) Extraction of endogenous MeCP2 from HEK293 nuclei **E)** (Left) FRAP recovery curves of MeCP2-GFP in NIH/3T3 cells. (Right) Times of half recovery, calculated using EasyFRAP. All figure graph data represent mean ± SEM and were analysed by one-way ANOVA followed by Tukey’s Multiple comparison test; ** p < 0.01, *** p < 0.001.

*The MeCP2 RBD binds RNA in cells and* in vitro

Given the unknown RNA specificity of the MeCP2 RBD, we assessed general MeCP2-RNA binding in HEK293 cells via biphasic separation and enrichment of protein crosslinked to bound RNA (**Fig. 3A**) (25, 26) which has successfully been used by others to verify RNA binding proteins (36, 37). The low mobility MeCP2-RNA species signal was reduced by MeCP2^ΔRBD^ compared to WT in the enriched samples, similar to positive and negative RNA Binding Protein (RBP) controls, FKBP25^WT^ and FKBP25^K22/23M^, respectively (**Fig. S4A**) (38). Urdaneta et. al showed that electrophoretic mobility of crosslinked protein-RNA complexes was dependent on RNA size (25); therefore, the reduction in the high molecular weight species indicates that the RBD may play a role in binding relatively large RNA molecules, whereas other MeCP2 modules are able to facilitate binding to smaller RNAs. The DNA binding protein Histone H4 was not captured in the enriched phase compared to input. RNase A treatment prior to enrichment eliminated the presence of RNA-bound protein, confirming that the enriched fractions are composed of protein bound to RNA.

**Figure 3.**
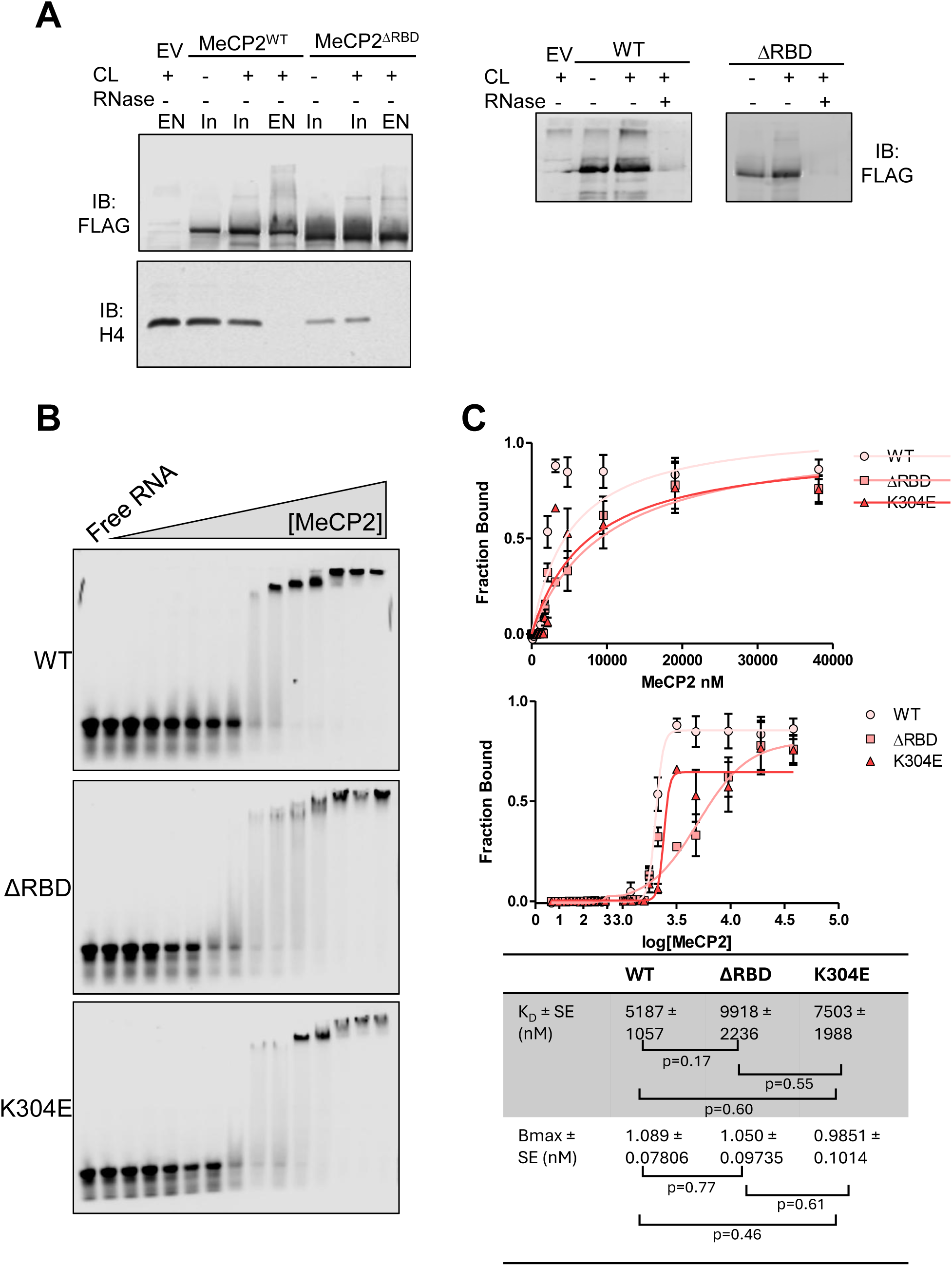
MeCP2’s NID/RBD facilitates RNA binding in cells and *in vitro*. **A)** Phenol-Toluene Extraction (PTex) of protein UV-crosslinked to RNA which appear in EN fractions from transiently transfected HEK293 cells. Top left: Flag-tagged WT and ΔRBD MeCP2; right bottom: endogenous Histone H4; Right panels: EN fractions of WT and ΔRBD MeCP2 PTex with and without RNase A treatment. Images represent 3-4 independent experiments. EV: Empty Vector; CL: Crosslink; In: Input. **B)** RNA Electrophoretic Mobility Shift Assays (REMSAs) with 100nM TYE665-labelled 21-mer siRNA probe corresponding to the human *NUP153* mRNA nucleotides 2297-2315 with 0.0, 0.3, 0.4, 0.7, 0.8, 1.0, 1.2, 1.5, 1.8, 2.1, 3.2, 4.8, 9.5, 19.1, and 38.1 µM protein as indicated. **C)** REMSA analysis. Top: Non-linear regression analysis of densitometric EMSA measurements (see Methods). Bottom: densitometric EMSA measurements where X values were transformed using X=Log_10_(X). Kd and Bmax p-values were acquired by Tukey multiple comparison calculation using the mean, standard error and N values.

To further validate NID/RBD binding to RNA, we performed RNA Electrophoretic Mobility Shift Assays (REMSAs) with recombinantly expressed full length MeCP2 (**Fig. S4B**). Lacking insight on RNA specificity from the non-canonical RBD of MeCP2, we performed EMSAs with a 5’-TYE665 fluorescently tagged dsRNA matching the 21-mer siRNA sequence previously identified as an MeCP2 binding partner (**Fig. 3B)**, but the affinity or domain specificity was not characterized at that time (5). Whereas MeCP2^WT^ had an apparent K_D_ of 5.2 µM ± 1.1 µM, the observed affinities of the ΔRBD and K304E mutants were reduced, with apparent K_D_s of 10.0 µM ± 2.2 µM and 7.5 µM ± 2.0 µM, respectively (**Fig. 3C**). The quantitative data are fairly noisy, making a definitive stance on the true difference in binding difficult to make, however there is room for some qualitative discussion. The observed smearing of MeCP2^ΔRBD^ indicates weaker, more transient binding to the probe compared to WT and K304E, resulting in a more gradual slope in the binding curve in **Figure 3C** (middle panel). While RNA binding to this particular probe is not completely ablated by mutation of the RBD, these data indicate that the NID/RBD region of MeCP2 contributes to dsRNA interaction *in vitro*. Whether other RNA species would provide clearer differences with stronger statistical strength between WT MeCP2 and NID/RBD mutants will be worth investigation in future studies.

During the preparation of this manuscript, *in vitro* data was published showing high affinity MeCP2 binding to C-rich RNA containing large stem-loops (6). Intriguingly, we noticed during our REMSA optimizations that addition of tRNA appears to compete with WT MeCP2 (**Fig. S4C)**, and thus we chose BSA as the non-competitive inhibitor.

Given that tRNA’s structure is comprised of multiple stem-loops, we questioned whether this tRNA competition is due to high affinity binding at the NID/RBD. Removal of the RBD, however, does not preclude tRNA’s competition with the siRNA probe, suggesting that tRNA binding to MeCP2 occurs outside the NID/RBD (**Fig. S4D**). MeCP2 RNA binding is thus likely modular and the NID/RBD facilitates RNA interaction but is not the only region of the protein capable of binding RNA. Taken together, these data show the non-canonical RBD of MeCP2 facilitates, but is not fully responsible for, RNA binding *in vivo* and *in vitro*. The other module(s) responsible for RNA binding, the specificity of them separately and combined, and whether RNA binding to domains outside the NID/RBD is biologically relevant will be important to tease out in future studies.

### MeCP2 binds to specific regions of the lncRNA NEAT1L

Non-canonical RBDs are often bound by regulatory non-coding RNAs. Evidence for MeCP2 interaction with long non-coding RNAs (lncRNA) exists; of the known MeCP2-lncRNA partners mentioned in the introduction, *NEAT1_2* is expressed in HEK293 cells (39, 40). Native RNA Immunoprecipitation (RIP) of MeCP2-FLAG from HEK293 isolated nuclei (41) shows marked decreases in *NEAT1_2* bound by ΔRBD and K316E MeCP2 compared to WT (**Fig. 4A**), similar to those seen in our REMSAs (**Fig. 3B, C**).

**Figure 4.**
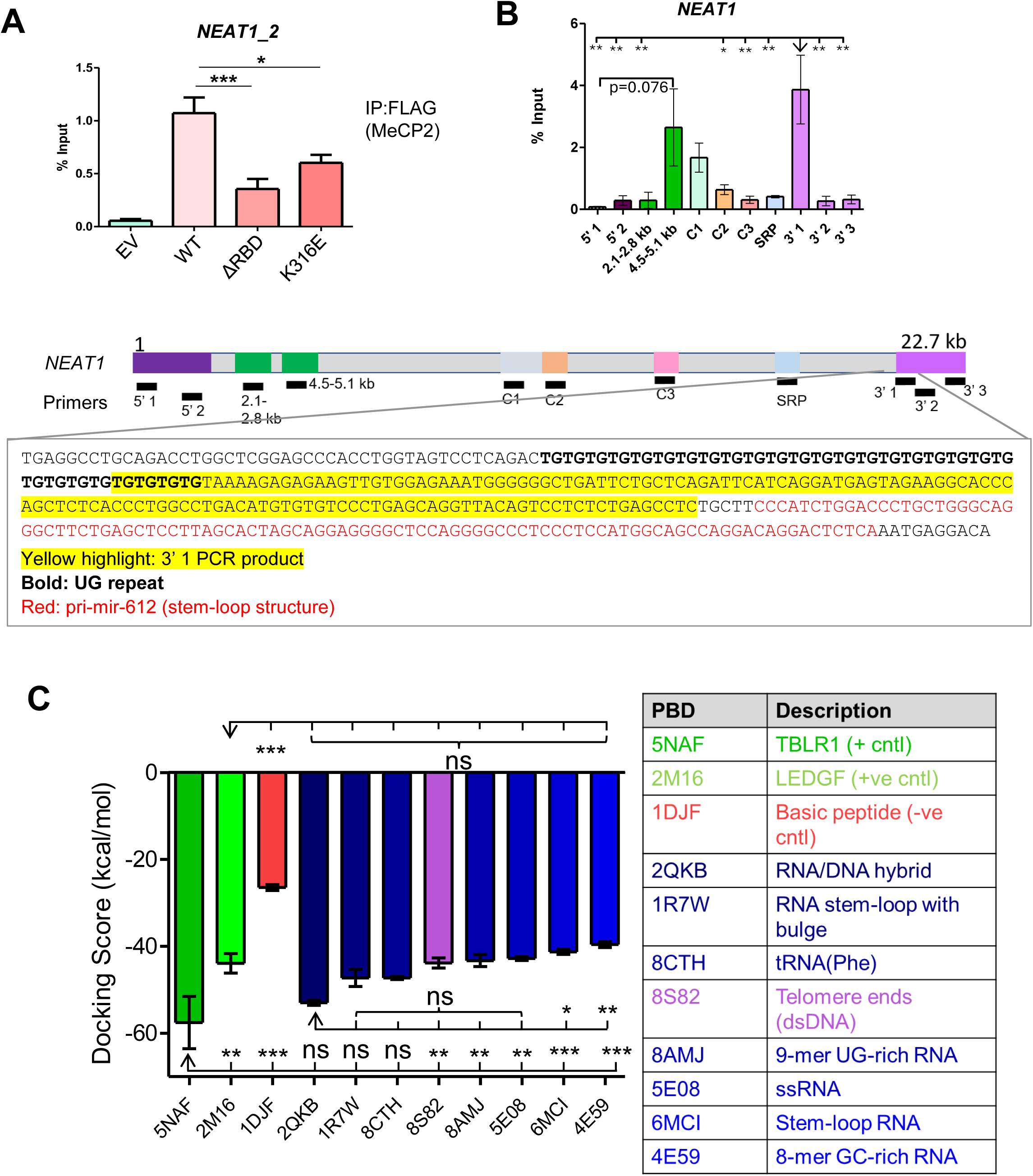
MeCP2 binds to specific regions of the lncRNA *NEAT1L*. **A)** Native RNA Immunoprecipitation (RIP) of *NEAT1_2* by FLAG-tagged MeCP2 from isolated nuclei of transfected HEK293 cells. **B)** Crosslinking WT MeCP2 RNA Immunoprecipitation RT-qPCR at selected regions of the *NEAT1_2* transcript after partial RNase digest (Left) and sequence/ structure features of interest from a part of the transcript enriched by WT MeCP2-FLAG (Below). Size range of partially digested RNA is indicated by agarose gel (Right). qPCR bar graphs represent mean + SEM, and one-way ANOVA with Tukey Multiple Comparison Test post-hoc test from 2-3 separate transfections and 3-4 technical replicates for each qPCR reaction. **C)** Molecular docking scores between the MeCP2 NID (PDB: 5NAF chain E) and other selected molecules. Data represent mean + SEM, and one-way ANOVA with Tukey Multiple Comparison Test; * p < 0.05, ** p < 0.01, *** p < 0.001.

**Figure 5.**
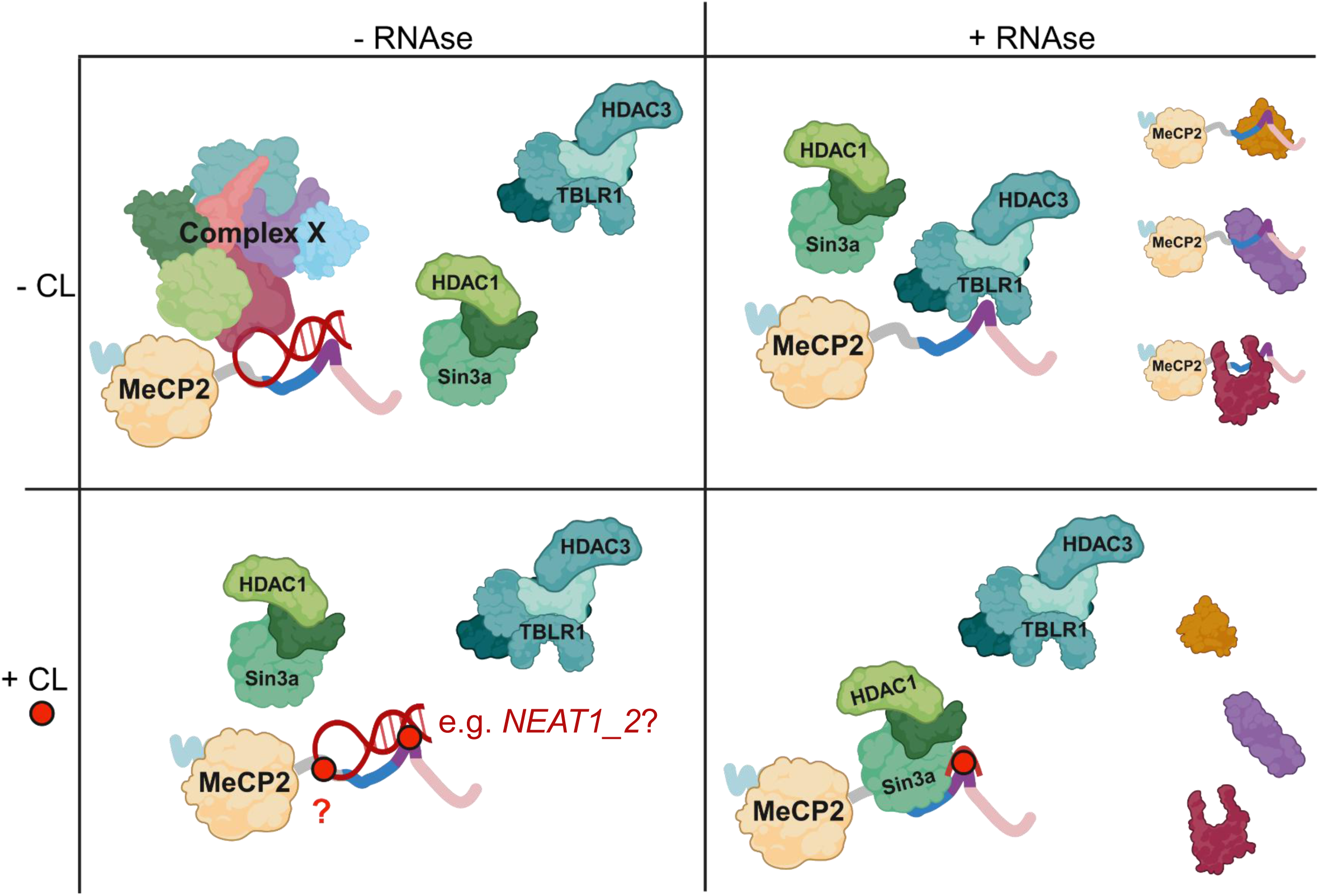
Schematic model for possible MeCP2 regulation by RNA at the RBD. In the presence of RNA, binding to certain proteins, such as TBLR1 and Sin3a, is limited, however there are other unknown protein complexes to which MeCP2 interacts through, perhaps, scaffolding RNAs that may bind at the RBD, at some other region of the protein, or both. When RNA is degraded, some large complexes disassociate from MeCP2, as indicated from sucrose density ultracentrifugation, but MeCP2’s total protein interactome increases through interactions at the RBD, such as to TBLR1, as shown by IP data. When UV crosslinking is applied to create a covalent link between RNA bound to MeCP2, RNase treatment does not result in the increased interactome or TBLR1 binding as before, but instead obstructs TBLR1 binding, allowing other proteins, like Sin3a, to bind MeCP2 at a domain outside the RBD.

To refine upon the specificity of MeCP2 to RNA, partially digested UV-crosslinked RNA was immunoprecipitated by MeCP2 to allow for qRT-PCR-based foot printing (42). Primers were designed by taking advantage of known functional *NEAT1_2* domains (43, 44), rather than examining the entirety of the 22 kb lncRNA (**Fig. 4B**). MeCP2 more effectively immunoprecipitated the 3’ and 5’ regions of *NEAT1_2*, which corresponds well with the known structure of the lncRNA, wherein *NEAT1_2* folds inward upon itself and self-associates into paraspeckles, where the inner core binds to constitutive paraspeckle proteins and the outer shell, composed of the ends of the RNA, interact with variable auxiliary proteins (45–47). Examining the area around the 3’-1 amplicon – due to partial RNase digestion potentially allowing IP of surrounding regions up to 200 nt – shows multiple sequence and secondary structural elements as possible candidates for specific binding, including a long stretch of UG-repeats as well as a stem-loop structure known to mimic a premature miRNA previously shown to sequester the microprocessor complex (47) (**Fig. 4B**). *In silico* molecular docking of the NID peptide crystal structure (PDB: 5NAF) using Molecular Operating Environment (MOE) software with various RNA and DNA species for which there were high quality crystal structures showed relatively similar binding preference for all the nucleic acid molecules as well as the known binding partner LEDGF positive control (PDB: 2M16) (48), although the docking score for NID-TBLR1 was highest of all molecules tested (**Fig. 4C**). An RNA/DNA hybrid had the highest docking score and was the only molecule with a non-significant difference compared to the TBLR1 positive control, thus representing a compelling candidate for further *in vitro* analysis.

## Discussion

With varying degrees of affinity for differently modified DNA as well as over 40 MeCP2-protein interactions guiding MeCP2 involvement in processes including transcriptional regulation, mRNA splicing, miRNA biogenesis and chromatin arrangement (1), it is clear that MeCP2 function is almost never black and white. Here, we add to these puzzling data by introducing the NID as an RBD that is important, but not wholly responsible, for RNA interaction. When Castello et. al (18) identified this region of the protein as an RNA binding module, they validated this finding in a fluorescence-based assay in cells. Perhaps because of the scientific field in which the paper was published, this finding has thus far gone relatively unnoticed by MeCP2-focused researchers, despite the known importance of this domain to RTT. The inability of MeCP2^WT^ to immunoprecipitate TBLR1 after crosslinking and RNase A treatment provided initial validation that regulation of MeCP2 interaction with the NCoR complex occurs through direct RNA interaction at the RBD. Mutation of the RBD reduced RNA binding in cells both in a global screen and to the lncRNA *NEAT1_2* compared to wild type as well as *in vitro* to a dsRNA probe to a similar degree as in the cellular data, about half. Reduced slow-moving MeCP2^ΔRBD^ concurrent with fast-moving species remaining in enriched PTex fractions agrees with the notion that the RBD is not the only RNA binding module in MeCP2. It also suggests that, while the *in vitro* data shows reduced MeCP2^ΔRBD^ binding to a 21-mer RNA oligonucleotide, in cells, the RBD may be more important as an accessory to bind to larger RNA molecules, such as lncRNAs like *NEAT1_2*, for example. More recently, MeCP2 was shown to recruit major satellite RNA to chromocenters in mouse cells through the TRD domain (49), indicating this as a compelling *in vivo* NID/RBD-RNA candidate target worth further study.

Seminal MeCP2 research showed that expression of a severely truncated form of MeCP2 comprised only of the MBD, the NID, and a splice acceptor into *Mecp2^-/y^* mice alleviated RTT phenotypes (23). At the time, the authors themselves acknowledged the paradox of two domains sufficiently conferring functionality to a protein known to be complex. This work alludes to a possible explanation for the seeming contradiction of the biologically functional minimal MeCP2. The overlapping TBLR1 and RNA binding domain, and our CoIP data showing negative regulation of MeCP2-TBLR1 binding by RNA indicates that MeCP2 interaction with these molecules is likely mutually exclusive. lncRNAs like *NEAT1_2* can dynamically regulate protein availability in the nucleus by sequestering them into paraspeckles (40, 50). *NEAT1* is downregulated in *MECP2*-KO neural progenitor cells, and induction of *NEAT1* alleviates dysregulated phenotypes (51), underscoring not only the importance of this lncRNA to RTT, but the complexity of the functional relationship between MeCP2 and its targets at the DNA and RNA level. The fact that the NID/RBD does not broadly change MeCP2-chromatin interaction speaks to the autonomous modularity of MeCP2 protein domains (52), which may be important for allowing MeCP2 to simultaneously interact with gene targets via the MBD and either co-repressor proteins or regulatory RNAs at the NID, for example.

Within the scope of this work, we were not able to delineate the sequence or structure specificity of the RBD. Our data indicate modularity of MeCP2-RNA interaction, so future studies will benefit from testing various RNA sequences with a range of MeCP2 single and double mutant proteins. Functional predictions of additional RBDs are difficult, owing to the intrinsic disorder of MeCP2, however, compelling evidence for other possible RNA binding modules in MeCP2 already exists. The first report of MeCP2 and other MBD proteins binding to RNA showed that the RG domain of Methyl CpG Binding Domain Protein 2 (MBD2) was responsible (5). RG/RGG domains are common in RNA binding proteins, with preference for G-quadruplexes or GC-rich dsRNA (53), although MeCP2’s RG domain is considerably smaller than that of MBD2. AT-hooks, of which MeCP2 has 3, have been shown to have some RNA recognition ability (54); one that is downstream the MBD and overlaps the RG domain of MeCP2 qualifies as what is called an extended AT-hook (eAT) (55, 56), which have preference for RNA over DNA, and long stem-loops over short hairpins, specifically. Finally, the recently published work by Melikishvili et al. (6) suggested the MBD as an RNA binding surface, although that was based on molecular docking simulations using the MBD alone (PDB: 6OGK) as the ligand as input.

This work invites more questions than those with which it began. The most pressing to answer will be to elucidate which RNAs are specifically bound *in vivo* and to identify which of those impact RTT pathology – by regulating MeCP2 binding to TBLR1 and other proteins, for example. The transcriptional regulator lens epithelium derived growth factor (LEDGF) was also recently shown to bind to MeCP2 at the NID (48), further expounding upon the functional minimal MeCP2 mystery as well as being an additional possible direct regulatory target of MeCP2 by RNA. MeCP2 is involved in mRNA splicing and microRNA biogenesis, where some research has shown RNA-dependent interaction with splicing proteins (11, 12); whether these RNA-dependent interactions occur at the NID/RBD requires further investigation. Finally, the effect of dynamic MeCP2 post-translational modifications (PTMs) on RNA binding will be an important addition to future work to clarify any complexity of MeCP2-RNA binding regulation. Brain activity causes MeCP2 to differentially bind DNA and TBLR1 in a PTM-dependent manner (57, 58) and PTMs have been shown to regulate RBP-RNA binding (59). There are known methylation, phosphorylation, acetylation and ubiquitination sites within the NID/RBD alone (56).

The data herein are foundational to understanding how MeCP2 is regulated by RNA at a pathologically relevant domain, knowing the full scope of which will be critical when considering direct and downstream effects of future MeCP2-directed therapies.

## Materials and Methods

### Plasmids and Cloning

Gene synthesis of the human WT and mutant *MECP2* E1 and E2 sequences were generated by Invitrogen, with the E1 and E2 sequences codon optimized for mammalian and bacterial expression, respectively. *MECP2E1* sequences were cloned into p3xFLAG-CMV^TM^-14 (Sigma) expression vector in-frame with the C-terminal FLAG-tag using 5’*Not*1 and 3’*Bam*H1 restriction sites. *MECP2E1* sequences were also cloned into pcDNA3.1 CT-GFP, in frame with GFP using 5’*Kpn*I and 3’*Xba*I restriction sites.

*MECP2E2* sequences were cloned in house by Q5 PCR amplification from pTYB1 vectors containing the codon optimized *MECP2E2* genes synthesized by Invitrogen and cloned into a pTEV6xHis-T7 vector using 5’*Nco*I and 3’*Eco*RI restriction sites. In-house cloning primers (5’-3’) were FWD: TAAGCACCATGGTTGCAGGTATGCTG, RVS: CATTGAGAATTCTTAGCTAACACGTTCGGTAAC

### Cell Culture

Cells were maintained at 37°C at 5% CO_2_ atmosphere with the following media components:

*HEK293, NIH/3T3, C2C12*: DMEM, 10% FBS, 1X PenStrep; *P19*: AMEM, 10% FBS, 1X PenStrep; *SHSY5Y*: DMEM/F12, 10% FBS, 1X PenStrep; *SKNSH*: EMEM, 10% FBS, 1X PenStrep.

### Cell Transfections

For the PTex and some of the HEK293 Co-IP transfections, the calcium phosphate method was used. Briefly, for each 10cm culture plate, cells were grown to approximately 75% confluence, then a 15 mL falcon tube was prepared with the following: 750 µl sterile distilled water (sdH_2_O), 20 µg DNA in TE buffer, and 55 µl freshly made 2.5M 0.2 µM-filtered CaCl_2_. 750 µl 0.2 µM-filtered 2x HEPES Buffered Saline (HBS) (280 mM NaCl, 1.5 mM Na_2_HPO_4_, 100mM HEPES pH 7; brought to pH 7.12) was added slowly, dropwise, to each falcon tube, while vortexing, then incubated at room temperature for 15 minutes. The DNA-Ca^2+^ complex solution was then added dropwise to cell plates, being careful not to disturb the cell layer. Cells were incubated for 14-16 hours, then transfection media was replaced with fresh complete media and cells were given 24 hours to recover and grow before harvesting. Otherwise, Lipofectamine 3000 Reagent (ThermoFisher) was used, following manufacturer instructions.

### Nuclear Isolation from Cells and Tissues

Cell culture: Pelleted cells were homogenized in 4 volumes of lysis Buffer A (0.25 M Sucrose, 60 mM KCl, 10 mM MES pH 6.5, 5 mM MgCl_2_, 1 mM CaCl_2_, 0.5% Triton X-100 and Roche Complete Protease Inhibitor without EDTA) by gentle pipetting and flicking, incubated on ice for no more than 5 minutes and centrifuged at 500 x *g* at 4°C for 10 minutes. The supernatant was discarded, and the pellet was resuspended in 2 volumes of nuclei suspension Buffer B (50 mM NaCl, 10 mM PIPES pH 6.8, 5 mM MgCl_2_, 1 mM CaCl_2_, and Roche Complete Protease Inhibitor without EDTA), or whichever volume brought the chromatin concentration to 1.5-2 mg/mL, based on A_260_ readings.

Adult mouse brain and liver: whole brain or liver was dounce-homogenized in 6 (brain) or 4 (liver) volumes of Buffer A and incubated on ice for 10 minutes then centrifuged at 600 x *g*, 4°C for 10 minutes. The supernatant was removed, and the pellet was resuspended in 12 (brain) or 8 (liver) volumes of Buffer A and immediately centrifuged as before for 5-8 minutes. The supernatant was removed, and the pellet was resuspended in 4 volumes of Buffer B using cut pipette tips and centrifuged as before for 10 minutes. The final pellet was resuspended in 2 volumes of Buffer B or whichever volume brought the chromatin concentration 1.5-2 mg/mL, based on A_260_ readings in a 0.5% SDS solution.

*Micrococcal Nuclease (MNase) Digestion-based Chromatin Fractionation (S1, SE, P)* Nuclei suspended in Buffer B (see *Nuclei Isolation*) were digested by Micrococcal nuclease (MNase) at 30 units/mg DNA for exactly 15 minutes in a shaking 37°C water bath. Some input sample was aliquoted before digestion. The digestion was stopped by adding EDTA to a final concentration of 20 mM on ice. The digest mixture was centrifuged at 9600 x *g* for 10 minutes at 4°C. The supernatant, S1, was transferred to a separate 1.5 mL microfuge tube and its A_260_ was taken before snap freezing. The pellet was harshly resuspended in 0.25 mM EDTA at an equal volume of the nuclei in Buffer B before digestion. The sample was then vortexed several times and tumbled at 4°C for 1 hour. Samples were then centrifuged at maximum speed for 10 minutes at 4°C. The supernatant, SE, was collected and its A_260_ was taken before snap freezing. Remaining pellets were dissolved in at least 3 volumes of 1X SDS sample buffer (125 mM Tris–HCl pH 6.8, 2% SDS, 20% glycerol, 1.43 M β-mercaptoethanol, and 0.2% bromophenol blue). Input samples were diluted in 4 volumes of sample buffer. Input and pellet SDS samples were sonicated in 30 second intervals with 30 second breaks on ice for 5-6 minutes at 60-80% power output with a Microson Ultrasonic Cell Disruptor MS-50 (Heat Systems Ultrasonics, Inc) and all SDS samples were boiled at 95°C for 3 minutes before SDS-PAGE gel loading.

### Co-localization Image Acquisition and Analysis (Confocal Microscopy)

C2C12 cells were seeded onto coverslips pre-coated at room temperature for at least one hour with 100 µg/mL Poly-D-Lysine in sterile distilled water and transfected with GFP constructs listed in the main text. Cells were then rinsed with PBS then fixed with ice-cold (pre-cooled at −20°C) 100% methanol at −20°C for 15 minutes. Cells were rinsed again in PBS and DAPI stained with NucBlue Fixed Cell ReadyProbes Reagent (ThermoFisher) according to manufacturer instructions. Coverslips were then mounted onto cleaned microscope slides with Dako Mounting Medium (Agilent Technologies). Images were acquired on an Olympus FV10-ASW Biological Confocal Laser Scanning Microscope and FluoView-1000 software. Co-localization analysis was performed by calculating Pearson’s Correlation Coefficient (PCC) using the JaCoP plugin in ImageJ. Statistical significance was tested with one-way analysis of variance (ANOVA) and Tukey’s post-test. Data represent 6-7 images from each of three biological replicates.

### Salt Extraction of Chromatin-Bound Protein

Solutions of 10 mM Tris (pH 7.50), 1 mM EDTA (pH 8.00) with increasing NaCl (0.4, 0.6, 0.7, 0.8, 0.9, and 1 M) were prepared. Equal amounts of the salt solutions were added to equal aliquots of nuclei preparations (see “*Nuclear Isolation from Cells and Tissues*”), vortexed well, and incubated on ice for 20 minutes. Samples were then centrifuged at 4°C for 10 minutes at 16.1 x 1000 *g*. The supernatant was removed from the pellet and appropriate volumes of SDS-PAGE sample buffer was added. Samples were stored at - 80°C.

### Co-Immunoprecipitation

Cell lines listed in the main text were first transfected as described above with FLAG-tagged MeCP2. For Protein Co-IPs, transfected cells in 10cm culture dishes were either UV-crosslinked at 150mJ/cm^2^ in a UV Stratalinker 2400 (Stratagene) on ice or immediately processed without crosslinking. FLAG-tagged proteins were immunoprecipitated using Sigma ANTI-FLAG M2 Affinity Gel according to manufacturer instructions, with or without the addition of 100 µg RNase A prior to separation of Input and incubation with FLAG M2 beads.

RNA Immunoprecipitations were performed as described in detail by our lab previously (41).

### Silver Stain

Samples run through 15% SDS polyacrylamide (37.5:1 polyacrylamide: bisacrylamide) gels were immediately incubated at room temperature with shaking in 1:1 5% acetic acid: 100% methanol for 30 minutes and washed twice in dH_2_O for 2 minutes each. A third dH_2_O wash was left overnight to remove yellow background. Gels were then sensitized in 0.02% sodium thiosulfate for 2 minutes and washed twice in dH_2_O for 30 seconds each time. Gels were incubated in 0.1% silver nitrate (w/v) for 30 minutes followed by another two 30 second dH_2_O washes. Gels were developed in 0.01% formaldehyde (v/v) diluted in 2% sodium carbonate (w/v) to a point that a yellow tint was detectable, at which point the formaldehyde/ sodium carbonate solution was replaced with fresh solution and allowed to fully develop (about 10 minutes). Gels were then washed and finally stored in 1% acetic acid. Images were acquired on a GelDoc XR+ system (BioRad).

### Western Blot

Polyacrylamide gels were transferred onto 0.2 μm nitrocellulose (Bio-Rad) or PVDF (Bio-Rad) membrane for 2-3 hours at 400mA on ice or 16 hours at 4°C at 30V/90mA in 40 mM: 192 mM tris: glycine plus 20% methanol. Membranes were blocked in 3-5% skim milk in PBS-Tween 0.1% for 1 hour at room temperature with shaking. Primary antibody incubation was done overnight at 4°C or 2 hours at room temperature. Primary antibodies used were: MeCP2 (M9317, Sigma®, St. Louis, MO, USA), H4 (rabbit serum produced in-house), Sin3a (ProteinTech 14638-1-AP), TBLR1 (Bethyl, A300-408A), HDAC1 (Santa Cruz Biotechnology, sc-81598). Secondary antibody incubation was carried out at room temperature for 1 hour, with three 10-minute PBS-Tween 0.1% washes before and afterwards. Secondary antibodies used were: ECL Anti-rabbit IgG (NA934, GE Healthcare), Donkey Anti-Sheep/Goat IgG (AB324P, EMD Millipore), IRDye® 800 Anti-rabbit IgG (611-132-122, Rockland Antibodies & Assays), IRDye® 680LT Anti-mouse IgG (926-68020, LI-COR). The latter two secondary incubations were carried out in the dark. Western blots were analysed using Li-Cor C-digit or BioRad GelDoc XR+ for chemiluminescent imaging or Li-Cor Odyssey (LI-COR Biosciences) for near-infrared imaging. Images were analyzed using Li-Cor Image Studio Version 5.2 software or ImageJ.

### Bacterial Protein Expression and Purification

BL21(DE3) cells (NEB) were transformed with bacterial plasmids described above. Fresh transformants were inoculated into LB broth containing 100 ug/mL Ampicillin (LB-Amp) then incubated at 37°C 250 RPM until the OD600 reached 0.5. Protein expression was induced by adding IPTG to a final concentration of 0.5 mM, and the culture was maintained at 25°C 250 RPM overnight. Cells were pelleted at 800 x *g* then snap frozen prior to purification.

Cell pellets were thawed on ice before resuspension in Lysis Buffer (50 mM NaH_2_PO_4_, 200 mM NaCl, 10 mM Imidazole, 10 mM 2-mercaptoethanol, 0.05% (w/v) CHAPS, 10% Glycerol, 1 mM PMSF, pH 8.0) at a 50X concentration factor per cell pellet wet weight. Lysozyme at a final concentration of 1 mg/mL was then added and incubated on ice for 30 minutes. Lysates were then sonicated 6 times for 30 seconds on ice, with intervening 30 second cooling periods. Lysates were then centrifuged at 10,000 x *g* for 30 minutes at 4°C. Lysate supernatant was added to Ni-NTA resin (Qiagen) equilibrated in lysis buffer and incubated with regular inversion at 4°C for 90 minutes. The Ni-NTA-lysate mixture was then washed in 20 column volumes of Wash Buffer 1 (50 mM NaH_2_PO_4_, 200 mM NaCl, 20 mM Imidazole, 0.05% (w/v) CHAPS, 10% Glycerol, 1 mM PMSF, pH 8.0), then 20 column volumes of Wash Buffer 2 (50 mM NaH_2_PO_4_, 500 mM NaCl, 50 mM Imidazole, 0.05% (w/v) CHAPS, 10% Glycerol, pH 8.0). Protein was serially eluted in Elution Buffer 1 (50 mM NaH_2_PO_4_, 500 mM NaCl, 500 mM Imidazole, 10% glycerol, 1 mM PMSF, pH 8.0) and Elution Buffer 2 (50 mM NaH_2_PO_4_, 500 mM NaCl, 750 mM Imidazole, 10% glycerol, 1 mM PMSF, pH 8.0). Samples were dialyzed in 10 kDa MWCO Slide-A-Lyzer dialysis cassettes (Cat. 66380 ThermoFisher) against 20 mM Sodium Phosphate pH 6.8, 150 mM NaCl and concentrated in 30kDa Amicon Ultra centrifugal filters (Millipore). Proteins were quantified and assessed for purity on Coomassie stained SDS-PAGE gels, using BSA as a standard.

### RNA Electrophoretic Mobility Shift Assay (REMSA)

Double stranded siRNA probe was purchased from IDT, with the sequence 5’ TYE665-ACCAAAUAAAACUGGCAAAUU. 1 µM RNA aliquots were heated at 95°C for 5 minutes in a heat block and allowed to cool to room temperature slowly overnight.

EMSAs were performed similar to (38), with some modifications. 10 µl reactions were prepared containing 20 mM Tris (pH 7.5), 150 mM potassium acetate, 1 mM EDTA, 10% glycerol, fresh 1 mM DTT and 0.1 mg/ml BSA as well as equal volumes of serially diluted protein in 25 mM Tris, 192 mM Glycine, 1 mM EDTA, pH 8.3 (TGE). Reactions were incubated on ice protected from light for 1 hour. 5% native polyacrylamide gels (37.5:1 polyacrylamide: bisacrylamide, 300 mM Tris-HCl pH 8.8, 0.1% (w/v) ammonium persulfate, 0.1% (w/v) TEMED) polymerized for 10 minutes then wells were rinsed and pre-run at 100V for 30-40 minutes in TGE at 4°C in the dark. Wells were rinsed again and 3 µl of each binding reaction was loaded into gels and run for 40 minutes. Gels were imaged in an ImageQuant 800 (Cytiva) and densitometric readings were measured using ImageStudio Lite software Version 5.2 (LiCor). Background measurements were first subtracted from Bound and Unbound signal measurements, then the Fraction of bound RNA was determined using the equation Bound/ (Bound + Unbound). These values were then plotted on a curve and used to calculate the apparent *K_D_* using the equation Y = Bmax*X/ (Kd + X), where Y is the fraction bound and Bmax is the maximum specific binding, in GraphPad Prism Version 5.01. Data represent 2-8 replicates from three independent experiments.

### Fluorescence Recovery After Photobleaching (FRAP)

NIH/3T3 cells were seeded and transfected at approximately 60-70% confluency on 35 mm MatTek glass bottom dishes. The Qiagen Effectene transfection kit was used with some modifications to the protocol. 100 µl of EC buffer was added along with 3 µl of Enhancer and 1 µg of DNA. This was incubated at room temperature for 20 minutes.

5 µl of Effectene was added, and the solution was incubated for another 20 minutes at room temperature before being added to the cells. Media was changed the following morning, and FRAP experiments were performed using a Zeiss LSM710 laser scanning confocal microscope with a 63x 1.4NA Oil DIC M27 Plan-Apochromat objective lens built around an inverted fully motorized Zeiss Axis Observer Z1 on 3 separate days per transfected construct. Entire heterochromatin domains (chromocenters) were bleached in nuclear mid-sections with a 488 Argon laser set to high power emission. 3 pre-bleach images were collected, and then after photobleaching time-lapse images were collected every 2 seconds. During imaging, cells were maintained in a live cell chamber at 37℃ with 5% CO2. FRAP curves were generated by registration of the time series using the rigid registration of the Fiji/ImageJ plug-in “Stack Reg”, subtracting background intensity and normalizing to the intensity of the whole nucleus throughout the FRAP experiment to account for the FRAP ROI and photobleaching of the entire cell (60, 61). As the result of repeated imaging (>30 observations per expression construct). T ½, Mobile and Immobile phase calculations were determined using the EasyFRAP tool (42).

### Phenol-Toluene Extraction (PTex)

Based on Urdaneta et al. (25). First, transfected HEK293 cells were washed twice in ice-cold sterile PBS then protein-RNA complexes were cross-linked by irradiation at 150 mJ/cm^2^ prior to harvesting by scraping in 600 µl PBS on ice. Non-irradiated controls were harvested immediately after PBS washes. From the 600 µl cell suspension in PBS, a 15 µl input sample was transferred to a new microfuge tube and snap frozen for later preparation. To the remaining cell solution, 200 µl each of neutral phenol (Sigma), Tolune (Sigma), and 1,3-bromochloropropane (BCP) (Sigma) was added. Samples were then mixed at 2000 RPM for 1 minute at 21°C in an Eppendorf thermomixer and subsequently centrifuged at 16,000 x *g* for 5 minutes at 4°C. The top aqueous layer (Aq1) was carefully transferred to a new 2 ml microfuge tube containing 300 µl solution D (5.85 M Guanidine isothiocyanate, 31.1 mM sodium citrate, 25.6 mM N-lauroyl sarcosine, 1% 2-mercaptoethanol). To RNase treated samples, 100 µg were added to Aq1 and incubated at 37°C for 10 minutes prior to solution D addition. To Aq1-Solution D, 600 µl neutral phenol and 200 µl BCP were added, and samples were mixed and centrifuged as before. The upper and lower three quarters of the second aqueous and organic layers (Aq2 & Org2), respectively, were removed and the following was added to the interphase (Int2): 400 µl H_2_O, 200 µl 95% EtOH, 400 µl neutral phenol, 200 µl BCP and samples were mixed and centrifuged as before. Aq3 and Org3 were carefully removed and Int3 was precipitated overnight in 9 volumes of 95% EtOH at −20°C. The next morning, samples were centrifuged at 16,000 x *g* for 30-35 minutes at 4°C. The supernatant was pipetted off and pellets were allowed to dry in a fume hood for no more than 20 minutes. Int3 pellets were finally solubilized in 30 µl 1 X SDS sample buffer, heated at 90°C for 5 minutes and snap frozen until use. Input samples were prepared by addition of 1 volume of lysis buffer (50 mM Tris-HCl, pH 7.4, 150 mM NaCl, 1 mM EDTA, 1% Triton).

### RT-qPCR

RNA from IP samples was extracted with TRIzol reagent (ThermoFisher) according to manufacturer instructions. cDNA synthesis was prepared with High-Capacity cDNA reverse transcription kit (Applied Biosystems). PCR samples were mixed with 0.5µM Forward and Reverse primers and SYBR Select Master Mix (ThermoFisher) and run on a MX3005P qPCR system (Stratagene) and MXPro software. Samples were run at least in triplicate. Data were analyzed by calculating 2 to the power of the ΔCt (between adjusted input and IP Ct averages) multiplied by 100%. Primers used are tabulated with references where applicable below and otherwise designed using the National Center for Biotechnology Information (NCBI) primer blast tool from the given accession code:

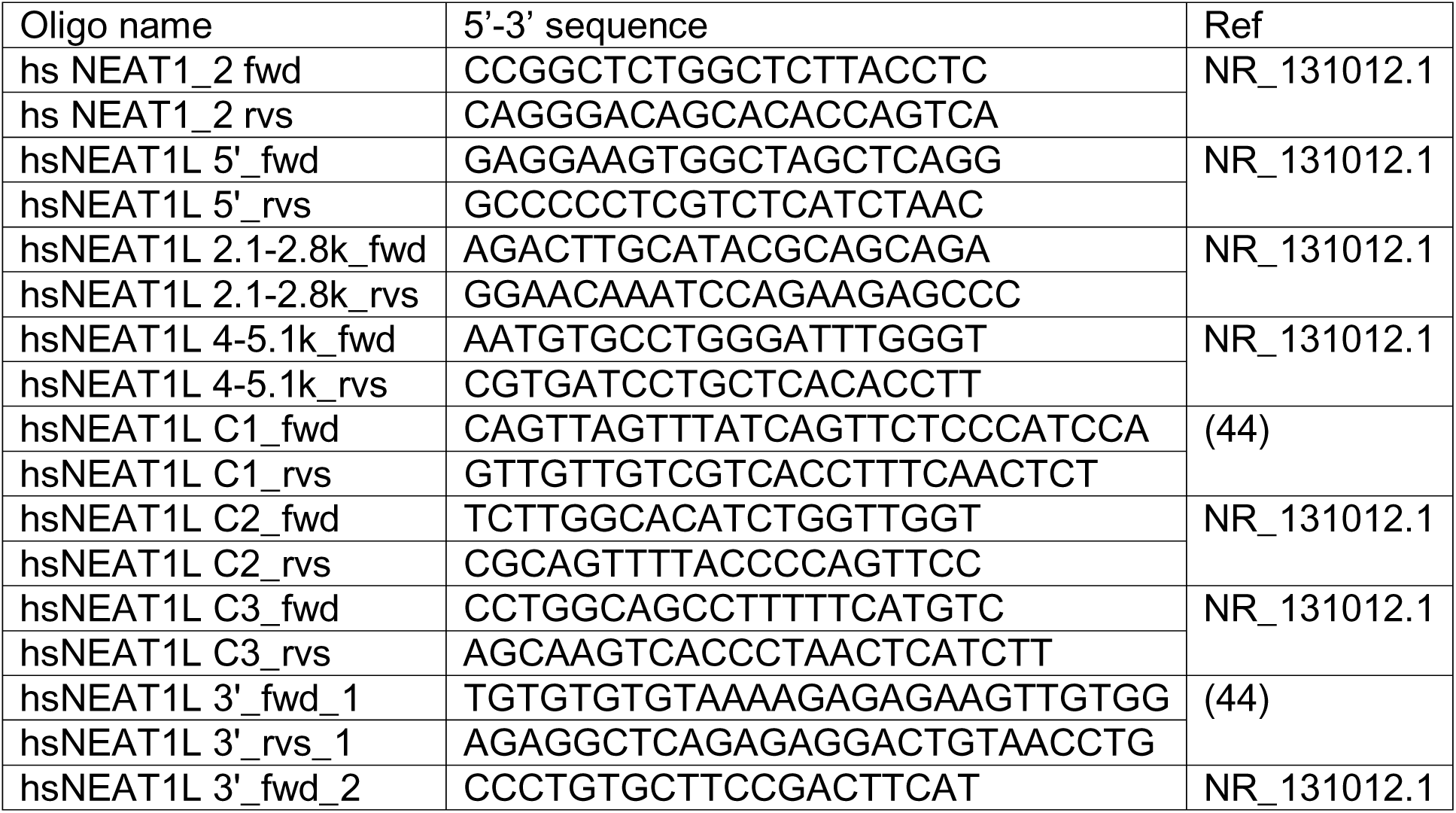
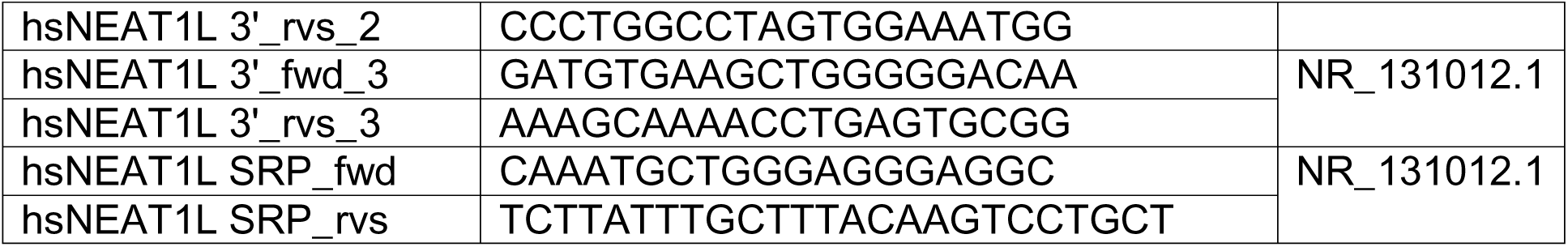

### MOE Molecular Docking

Molecular docking was performed as previously described (62). Briefly, the indicated PDB files were prepared in the Molecular Operating Environment (MOE) 2022.02 software by removing water and ligand molecules and adding hydrogen atoms using the Quick Prep tool. Molecule-Protein docking was performed with default parameters, and the top-scoring prediction was counted as one replicate, of which contacts were identified and recorded. Because of the small size of the NID peptide from the 5NAF crystal structure, it was categorized as the ligand, and the molecules as receptor. Three replicates were analysed by One-way ANOVA with Tukey’s Multiple Comparison Test using GraphPad Prism version 5.01.

### Sucrose gradient ultracentrifugation (RNA dependence)

We modified the protocol described in Caudron Herger et al., 2019 (24). Nuclei were prepared for HeLa S3 cells and absorbance at 260 nm was measured (See *Nuclei Isolation*). Nuclei were pelleted at 600 x *g* for 10 minutes at 4°C. The supernatant was removed, and nuclei were resuspended in 1 mL IP lysis buffer (50 mM Tris HCl, pH 7.4, 150 mM NaCl, 1 mM EDTA, 1% Triton) with complete protease inhibitor (Roche); RNase inhibitor was added only to control tubes. Nuclei were tumbled at 4°C for 2 hours. To ensure nuclear lysis, samples were frozen in liquid nitrogen and thawed at 37°C twice.

Microscopy using Trypan blue indicated degree of nuclear lysis. Nuclear lysates were cleared at 13,000 x *g* for 10 minutes at 4°C. Supernatant was transferred to a new tube. For nuclease treatment, 100 μg of RNase A was added to respective tubes and incubated at 37°C for 10 minutes, then kept on ice. About 50 μl of sample from each tube was aliquoted as input. Sample volume corresponding to 1 mg of chromatin was used for each gradient (about 1 mL). 34 mL 5-50% sucrose gradients were prepared by adding 17mL each of 5% sucrose and 50% sucrose in 100 mM NaCl, 10 mM Tris, pH 7.5, and 1 mM EDTA to the left and right wells, respectively, of a gradient maker. Tubing from the gradient maker into ultracentrifuge tubes were secured while making the gradient. Nuclear lysate supernatants were gently overlain onto gradients using a 200 μl cut pipette tip. Samples were centrifuged in a SW 32 Ti swinging bucket rotor at 30,500 RPM (114,000 x *g*) at 4°C for 19 hours. 35 1 mL fractions were collected using a Bio-Rad Model 2110 Fraction collector at 1 mL/minute. Glass tubing leading into ultracentrifuge tubes were very carefully inserted to the bottom of the tube and secured before fraction collection. Tubing was rinsed with Nuclease-free water and emptied before, between, and after fraction collection. 1 mL fractions were parafilmed and kept on ice until use. To prepare samples for western blot analysis, 300 μl of respective fractions was added to ∼9 cm pre-wet 3.5 kDa dialysis tubing. Samples were dialyzed in 4 L 4°C ddH_2_O for 6 hours, with stirring and water change at the 3-hour mark. Dialyzed samples were transferred to labeled 1.5 mL microfuge tubes with holes poked in the lid. Samples were flash frozen in liquid nitrogen and lyophilized overnight. Lyophilized samples were dissolved in 10 μl 1X SDS sample buffer and boiled at 95°C for 4-5 minutes. All 10 μl was loaded for each western blot.

## Acknowledgements

Thank you to Dr. Christopher Woodcock for the pTYB1 vector. A sincere thank you to Anna Mikhailov and Stephen Pastore for their help in the Vincent lab.

## Funding

This work was supported by the Centre for Addiction and Mental Health (CAMH) Foundation Discovery Fund grant to John Vincent and Juan Ausió.

## Conflict of Interest Statement

The Centre for Addiction and Mental Health (J.B.V) holds a patent and receives royalties related to diagnostic screening of *MECP2*. The authors otherwise declare no conflicts of interest.

## Author Contributions

K.G., H.S., M.H, and J.A. designed and conducted the FRAP experiment. T.M. and K.G. conceived and performed molecular docking simulations. K.G., C.J.N. and J.A. designed, and K.G. performed remaining experiments. K.G., C.J.N., J.A., and J.B.V. prepared the manuscript.

