## Supplemental Data for "MeCP2 NID interaction with RNA: Implications for Rett Syndrome-Relevant Protein Regulation"

### Supplementary Figures

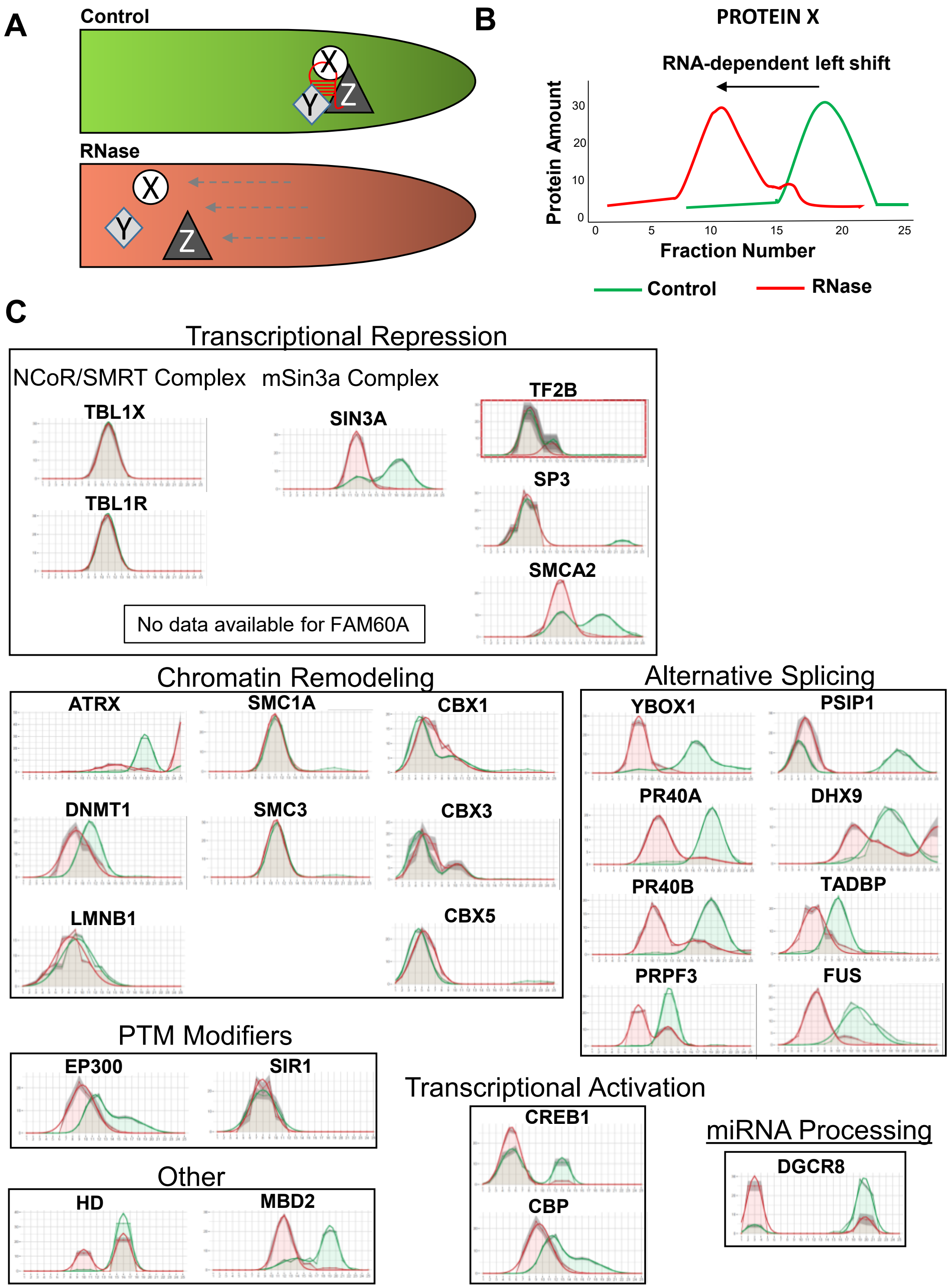


**Figure S1. R-Deep analysis of known MeCP2-protein partners.**

**A-B)** Illustrations of the process and data output for RNA-dependent protein complex disassembly after RNase treatment. **C)** RNA dependent protein complex plots of known MeCP2-interacting proteins generated from the R-Deep HeLa S3 cell database categorized based on the molecular process with which they are associated.

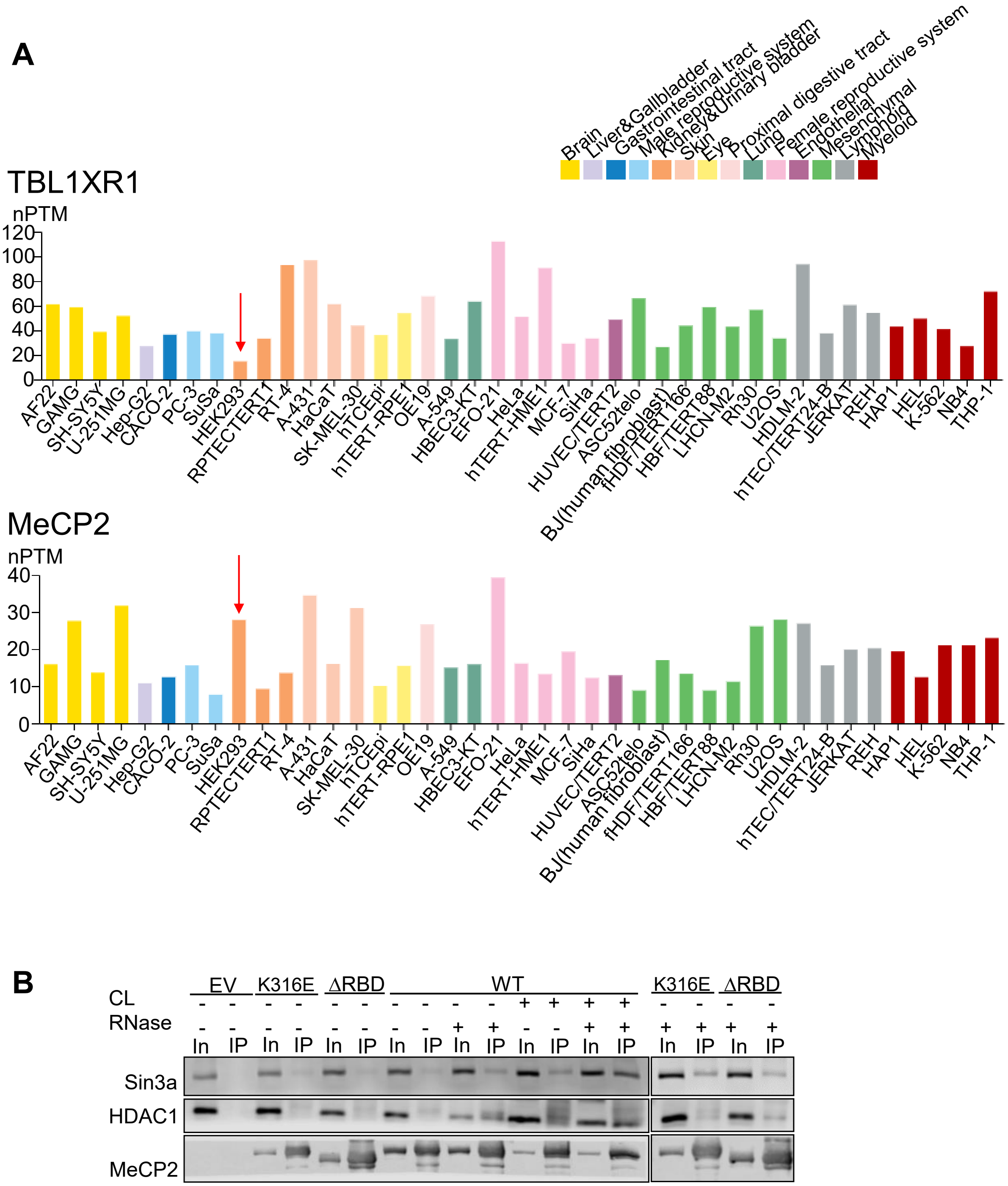


**Figure S2**. **Supplement to Figure 1 CoIP data.**

**A)** mRNA expression data indicates a relatively low ratio of *TBL1XR1* to *MECP2* in HEK293 cells. Protein atlas data of *TBL1XR1* mRNA expression profiles in different cell lines (SubCell tab). Red arrow denotes HEK293. Image credit: Human Protein Atlas, available from https://www.proteinatlas.org/humanproteome/subcellular. **B)** Co-IP and western blot of Sin3a and HDAC1 by transiently transfected Flag-tagged MeCP2 with indicated conditions. Images represent at least three separate experiments.

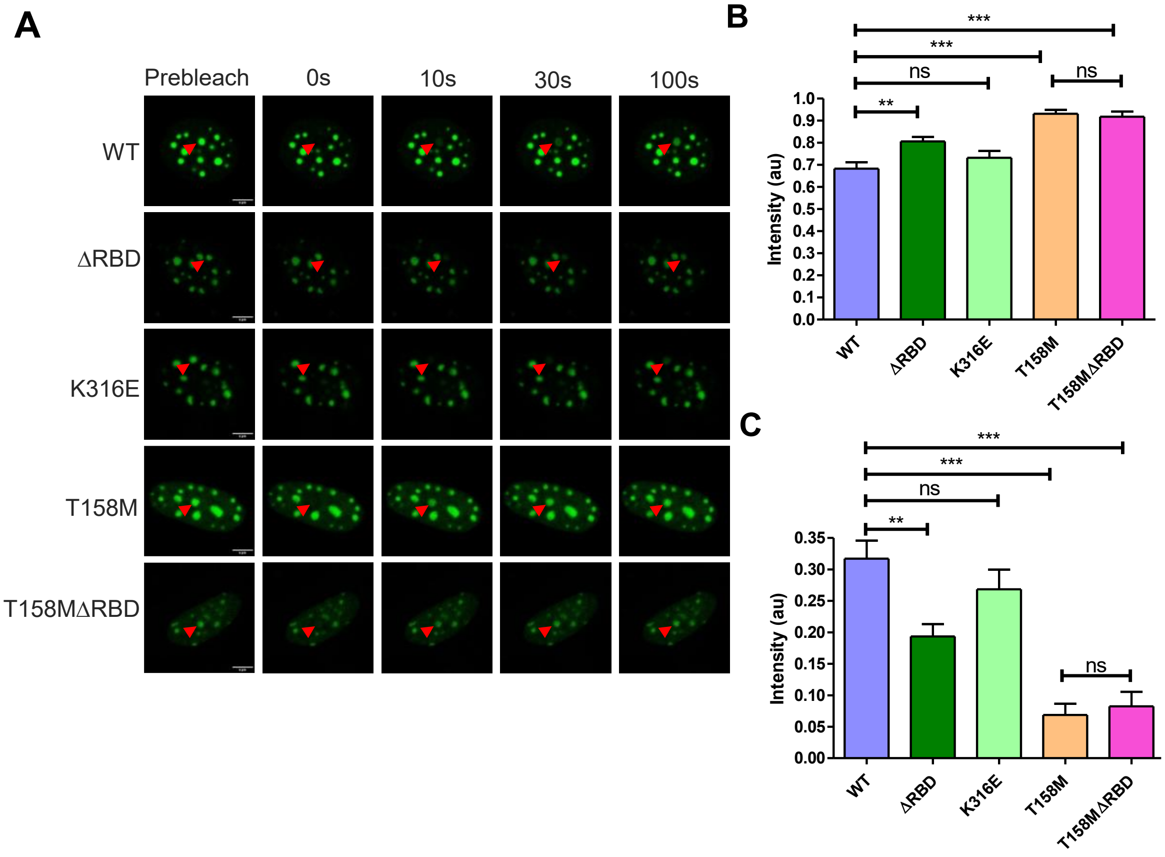


**Figure S3. Supplement to Figure 2 FRAP data.**

**A)** Representative time-stamp images of nuclei pre- and post- bleaching, with the bleached MeCP2-GFP-bound chromocenters indicated with red arrows. **B and C)** Mobile and Immobile phase calculations from FRAP data, calculated using the EasyFRAP tool and analyzed in GraphPad Prism. Data represent mean + SEM, and one-way ANOVA with Tukey Multiple Comparison Test; ** $p < 0.01$, *** $p < 0.001$,

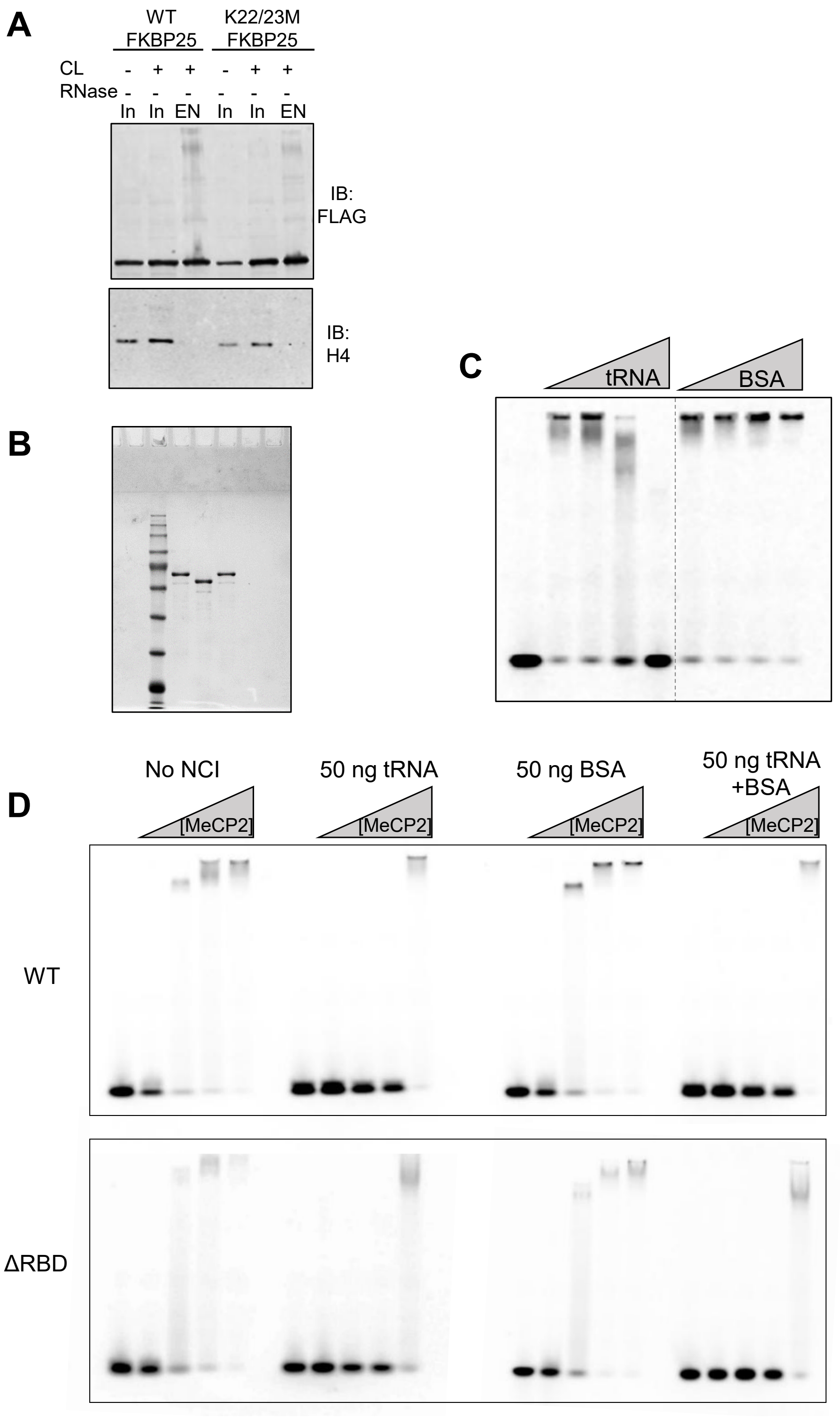


**Figure S4. Supplement to Figure 3 EMSAs.**

**A)** Phenol-Toluene Extraction (PTex) of protein UV-crosslinked to RNA which appear in EN fractions from transiently transfected HEK293 cells. Top: Flag-tagged WT and

K22/23M mutant FKBP25 (positive and negative RNA Binding Protein controls, respectively); bottom: endogenous Histone H4. **B)** Coomassie stained SDS-PAGE gel of recombinantly expressed and purified MeCP2$^{WT}$, $^{\Delta RBD}$, and $^{K304E}$. **C)** REMSAs with 100 nM TYE665-labelled 21-mer siRNA probe corresponding to the human *NUP153* mRNA nucleotides 2297-2315 with 19μM WT MeCP2 and 12.5, 25, 50, and 100 ng of yeast tRNA (left) or BSA (right). The first lane contains only RNA. **D)** REMSAs as in (C), but containing 50 ng of either tRNA, BSA, or both, and increasing MeCP2 concentrations from 0, 0.6, 2.4, 9.5, and 38.1 μM. NCI: Non-competitive inhibitor.
